# Evaluation of tools for long read RNA-seq splice-aware alignment

**DOI:** 10.1101/126656

**Authors:** Krešimir Križanović, Amina Echchiki, Julien Roux, Mile Šikić

## Abstract

**Motivation:** High–throughput sequencing has transformed the study of gene expression levels through RNA-seq, a technique that is now routinely used by various fields, such as genetic research or diagnostics. The advent of third generation sequencing technologies providing significantly longer reads opens up new possibilities. However, the high error rates common to these technologies set new bioinformatics challenges for the gapped alignment of reads to their genomic origin. In this study, we have explored how currently available RNA-seq splice-aware alignment tools cope with increased read lengths and error rates. All tested tools were initially developed for short NGS reads, but some have claimed support for long PacBio or even ONT MinION reads.

**Results:** The tools were tested on synthetic and real datasets from the PacBio and ONT MinION technologies, and both alignment quality and resource usage were compared across tools. The effect of error correction of long reads was explored, both using self-correction and correction with an external short reads dataset. A tool was developed for evaluating RNA-seq alignment results. This tool can be used to compare the alignment of simulated reads to their genomic origin, or to compare the alignment of real reads to a set of annotated transcripts.

Our tests show that while some RNA-seq aligners were unable to cope with long error-prone reads, others produced overall good results. We further show that alignment accuracy can be improved using error-corrected reads.

**Availability:** https://github.com/kkrizanovic/RNAseqEval

## 1 Introduction

Over the past ten years, the use of next generation sequencing (NGS) platforms, in particular Illumina, has expanded to dominate the genome and transcriptome sequencing market. Their sequencing-by-synthesis approach is indeed much cheaper and faster than the previously used Sanger sequencing. Recently, two new sequencing technologies, the so-called “third generation sequencing technologies”, have emerged, that produce longer reads and hold numerous promises for genomic and transcriptomic studies.

First, the single-molecule sequencing technology unveiled in 2010 by Pacific Biosciences (PacBio), produces reads up to a few tens of thousands of base pairs long. However, raw reads (“subreads”) display significantly higher error rate (∼10-20%) than reads from the Illumina technology (∼1%) (Schirmer *et al*., 2015; Ross *et al*., 2013; GLENN, 2011). To reduce error rates, circularized fragments are sequenced multiple times and the subreads produced can be reconciled to produce higher-quality consensus “Reads of Insert” (ROIs, previously called Circular Consensus Reads). However, there is a trade-off between the ROIs length and accuracy because, with longer fragments accumulating fewer sequencing passes.

Second, the portable MinION sequencer unveiled in 2014 by Oxford Nanopore Technologies (ONT), produces even longer reads (up to a few hundreds of thousand base pairs long), but with even higher error rates. Using the R7.3 chemistry, raw reads (“1D” reads) had an error-rate of more than 25%, while consensus “2D” reads (where template and complement of double-stranded fragments are successively sequenced and reconciled) displayed 12-20% error rate (Laver *et al*., 2015; Sović *et al*., 2016). It is likely that improvement in the chemistries (notably the recently released R9 and R9.4) has reduced error rates (http://lab.loman.net/2016/07/30/nanopore-r9-data-release).

For transcriptomic studies, long reads of these third generation sequencing technologies should be very helpful in the challenging task of identifying isoforms, and estimating reliably and precisely their abundances (Łabaj *et al*., 2011; Garber *et al*., 2011). It is unclear though whether high error rates will allow precise identification of exon-exon junctions required for proper discrimination of isoforms that are very similar in sequence (e.g., NAGNAG splicing).

The aim of this work was to determine whether currently available RNA-seq splice-aware aligners were able to handle third generation sequencing data, namely much longer read length and significantly higher error rate. Such a benchmark of RNA-seq alignment tools and pipelines, previously performed on both real and synthetic Illumina reads (Engström *et al*., 2013) proved to be very helpful for the community of end-users but, to our knowledge, was not yet performed on third generation sequencing data.

Splice-aware RNA-seq alignment tools can be divided into two groups. First, guided splice-aware aligners, use the genome sequence and known gene annotations to calculate gene or transcript abundance, but cannot be used to identify new splice junctions. Second, de novo splice-aware aligners can align RNA-seq reads to a reference genomic sequence without prior information on gene annotations.

In this paper, we chose to focus on de-novo splice aware aligners and on third generation sequencing data.

BBMap is to our knowledge the only tool claiming support of both PacBio and ONT reads (Bushnell *et al*., 2014). It uses short k-mers to align reads directly to the genome, spanning introns to find novel isoforms. It uses a custom affine-transform matrix to generate alignment scores.

A tutorial, developed by the PacBio team (available at https://github.com/PacificBiosciences/cDNA_primer/wiki/Aligner-tutorial:-GMAP,-STAR,-BLAT,-and-BLASR) recommends modified sets of parameters for the alignment of PacBio reads with STAR and GMAP, based on in-house testing. STAR (Dobin *et al*., 2013) employs sequential maximum mappable seed search in uncompressed suffix arrays followed by seed clustering and stitching procedure. It detects novel canonical, non-canonical splices junctions and chimeric-fusion sequences. GMap (Wu and Watanabe, 2005) is a part of GMAP/GSNAP package and uses dynamic programming to find an optimal global chain of short kmers.

In our tests we included TopHat2 (Kim *et al*., 2013), the most popular aligner for Illumina reads. TopHat2 implements a two-step approach where initial read alignments are first analyzed to discover exon-exon junctions, which are then used in the second step to determine the final alignment. HISAT2, the successor of Tophat2, was also included. It uses a global FM-index, as well as a large set of small FM-indexes (called local indexes) that collectively cover the whole genome. This strategy enables effective alignment of RNA-seq reads spanning multiple exons (Kim *et al*., 2015).

In the event that aligners are unable to cope with high error rates in the reads, we tested if the addition of an error-correction step before the mapping step could be useful. Recent tools have been developed that allow error correction of reads from third generation sequencing technologies, taking advantage of the redundancy within each dataset, or combining them with second generation sequencing datasets (Bradley *et al*., 2012). The latter (so-called “hybrid”) approach has already been used to obtain a comprehensive characterization of the transcriptome of the human embryonic stem cell (Au *et al*., 2013). In this work we applied both approaches and quickly discuss their merits.

## 2 Methods

Since the actual origin of reads in real datasets is unknown and can only be estimated through the alignment process, real datasets are not best suited to assess the performance of alignment tools. The accuracy and precision of aligners can be assessed on synthetic data, but in return simulators fail to mimic every aspect of real-life datasets, potentially biasing the benchmark results. In this work, we thus decided to use both simulated and real datasets.

All real datasets consist of RNA converted to cDNA and amplified prior to sequencing. For simulation, we have used the PacBio reads simulator PBSIM (Ono *et al*., 2013). Several datasets were simulated with different parameters, and using the annotated transcriptome of different organisms (the baker’s yeast *Saccharomyces cerevisiae*, the fruit fly *Drosophila melanogaster*, and human chromosome 19; see Supplementary data).

For the purpose of comparison, one ONT MinION dataset was also simulated using PBSIM, setting the parameters according to the statistics of ONT MinION R9 real data. While a PacBio simulator is not entirely appropriate for ONT MinION data, we felt that mimicking their read length and error profile (frequency of insertions, deletions and mismatches) should provide some useful insight. At the time of our simulation experiments, we were unaware of a dedicated MinION reads simulator. Since then, we became aware of NanoSim (Yang *et al*., 2017), but due to time constraints decided not include it in our benchmark.

In order to explore the effect of read error correction on alignment, the highest quality real PacBio dataset was error corrected using the recent consensus tool Racon (Vaser *et al*., 2017). Both correction using external Illumina reads and self-correction were explored.

The description of simulated datasets generation can be found in the Supplementary material. Table 1 shows relevant statistics of test datasets. As can be seen from the table, datasets vary in size and complexity. For example, datasets 2 and 4 have similar size because they were generated using the same approximation of the gene coverage histogram, however, since MinION ONT reads are on average longer than PacBio reads, dataset 2 contains more reads than dataset 4.

**Table 1.** Test dataset statistics

| Data set | Type | Organism | Technology | Size | No. genes | No. reads | % AS genes |
| --- | --- | --- | --- | --- | --- | --- | --- |
| 0 | Real | *D. melanogaster* | Illumina | 1 GB | NA | 4,000,000 | NA |
| 1 | Synthetic | *S. cerevisiae* | PacBio ROI | 400 MB | 6,000 | 185,000 | 0 |
| 2 | Synthetic | *D. melanogaster* | PacBio ROI | 1.4 GB | 7,000 | 412,000 | 10 |
| 3 | Synthetic | *Homo sapiens*, chr. 19 | PacBio ROI | 200 MB | 1,520 | 84,000 | 60 |
| 4 | Synthetic | *D. melanogaster* | ONT R9 2D | 1.4 GB | 7,000 | 342,000 | 10 |
| 5 | Real | *D. melanogaster* | PacBio ROI | 1 GB | NA | 192,000 | NA |
| 6 | Real | *D. melanogaster* | PacBio Subreads | 1 GB | NA | 243,000 | NA |
| 7 | Real | *D. melanogaster* | PacBio ROI error-corrected | 500 MB | NA | 192,000 | NA |
| 8 | Real | *D. melanogaster* | ONT R9 2D | 120 MB | NA | 40,000 | NA |

### 2.1 Datasets

To generate simulated datasets, we used PBSIM version 1.0.3, downloaded from https://code.google.com/archive/p/pbsim/.

Synthetic datasets were created from the following organisms:

- *Saccharomyces cerevisiae* S288 (baker’s yeast)
- *Drosophila melanogaster* r6 (fruit fly)
- *Homo Sapiens* GRCh38.p7 (human)

Reference genomes for all organisms were downloaded from http://www.ncbi.nlm.nih.gov.

PBSIM is intended to be used as a genomic reads simulator, taking as input a reference sequence and a set of simulation parameters (e.g., coverage, read length, error profile). To generate RNA-seq reads, PBSIM was applied to a set of transcripts generated from a particular genome using the gene annotations downloaded from https://genome.ucsc.edu/cgi-bin/hgTables. To make the datasets as realistic as possible, real datasets were analyzed and used to determine simulation parameters. Real gene expression datasets were used to select a set of transcripts for simulation (downloaded from http://bowtie-bio.sourceforge.net/recount/; core (human), nagalakshmi (yeast) and modencodefly (fruit fly) datasets ONT MinION were used).

- Dataset0 was randomly subsampled from available Illumina data (from 130 GB to 1GB) to obtain a coverage in the same order of magnitude as other datasets.
- Dataset1 was generated using *PBSIM* and an error profile obtained from PacBio ROI reads using *Saccharomyces cerevisiae* transcriptome.
- Dataset2 was generated using *PBSIM* and an error profile obtained from PacBio ROI reads using *Drosophila melanogaster* transcriptome.
- Dataset3 was generated using *PBSIM* and an error profile obtained from PacBio ROI reads using human chromosome 19 transcriptome. Only one human chromosome was used to keep the dataset size appropriate for the test.
- Dataset4 was generated using *PBSIM* and an error profile obtained from ONT MinION R9 reads using *Drosophila melanogaster* transcriptome.
- Dataset5 was randomly subsampled from available PacBio ROI reads.
- Dataset6 was randomly subsampled from available PacBio subreads.
- Dataset7 was obtained by error correcting available PacBio ROI reads using self-correction.
- Dataset8 was obtained by using all available ONT MinION R9 2d reads.

Real RNA-seq datasets used in this benchmark were generated from *D. melanogaster*. Technical replicates of the same sample were sequenced with three different technologies: Illumina HiSeq, PacBio RSII and ONT MinION. Illumina data were used for base-line comparison of all tested tools and for error correction of PacBio reads. PacBio and MinION data were used to assess aligners’ performances and to determine error profiles that were then used for simulation of synthetic data. In total we used:

- 1GB of Illumina reads, subsampled randomly from a larger dataset of 130GB. The reads were of size 101bp.
- Over 5GB of PacBio subreads, sequenced from 3 different size fractions of transcripts (1-2 kb, 2-3 kb and 3-7 kb, 2 SMRT-cells sequenced for each size fraction). This corresponded to about 2GB of Reads of Insert extracted from the subreads.
- 350MB of ONT MinION reads using the R9 chemistry. Because of the very low quality of 1D reads, only 2D reads were used in this benchmark.

### 2.2 Simulated data preparation

Simulated datasets were generated using the following workflow:

1. Analyze real datasets to determine error profiles.
2. Filter annotations (keep only primary assembly information) and unify chromosome names.
3. Separate annotations for genes with one isoform and genes with alternative splicing, keeping up to 3 isoforms randomly for each gene with alternative splicing.
4. Generate a transcriptome from processed annotations and a reference genome.
5. Analyze gene expression data and determine gene coverage histogram (Figure 1).
6. Approximate gene coverage histogram with 3 points to determine coverage and number of genes in simulated dataset (Figure 1). Scale coverages proportionally down to make a smaller dataset, more suitable for testing.
7. Extract 6 subsets of sequences from generated transcriptome, 3 for genes with single splicing and 3 for genes with alternative splicing. Each set contains a number of transcripts corresponding to the number of genes from a previous step.
8. Using *PBSIM*, simulate reads on each generated subset of transcriptome, using coverages determined in step 6 and error profiles determined in step 1.
9. Combine generated reads into a single generated dataset.

**Figure 1.**
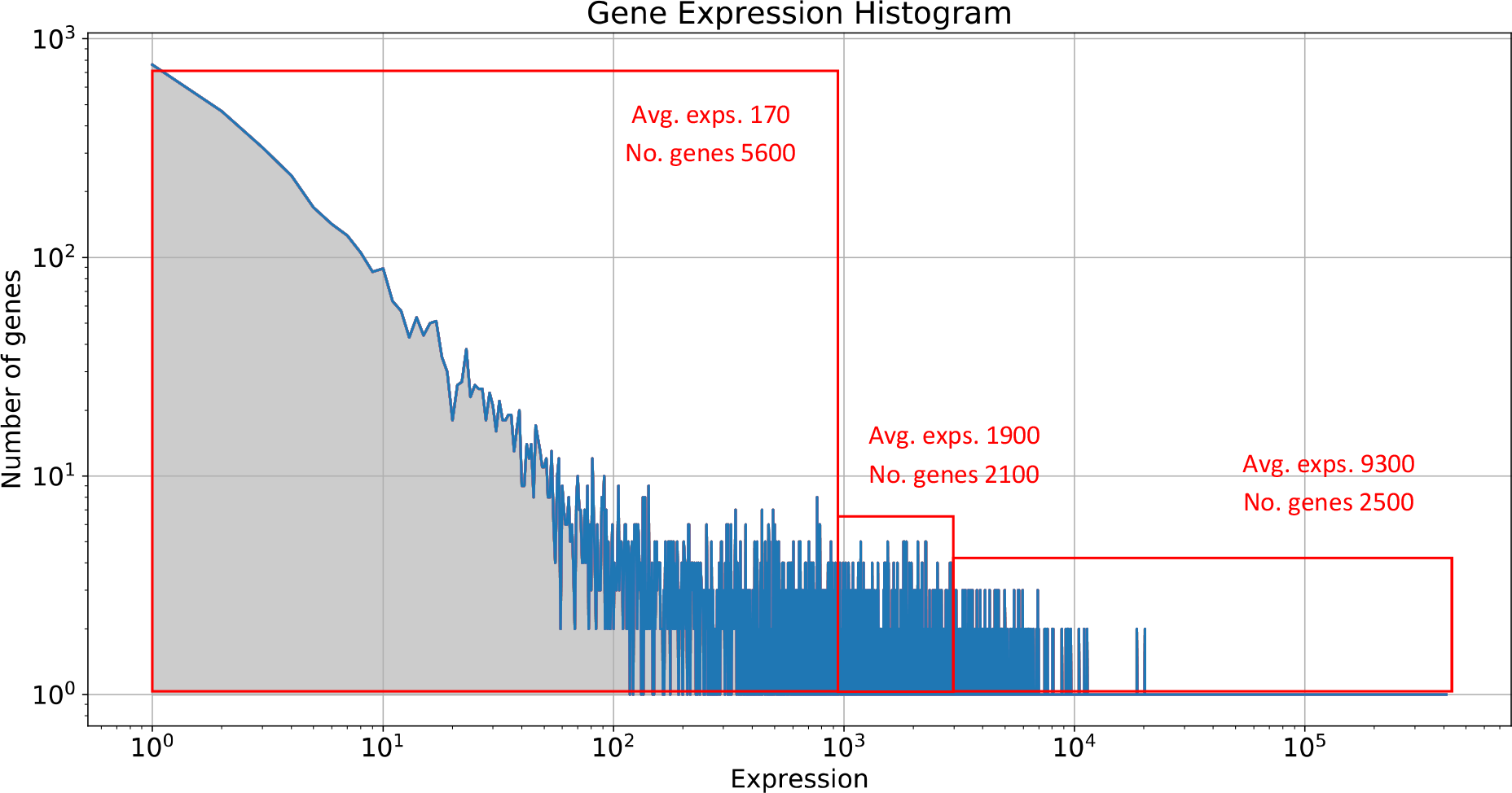
Data preparation step 6: Approximating gene expression with three points, applied to dataset 2. Three points were chosen as a compromise between achieving simple simulation and realistic datasets.

For simplicity, we rounded the coverage and number of genes from each transcriptome subset. For example, Table 2 shows the numbers used to generate dataset 2 (*D. melanogaster*). The annotation includes roughly 23,000 genes with a single isoform and 3,000 genes with alternative splicing. Rounding up the ratio, we have decided to simulate 1/10 genes with alternative splicing and 9/10 genes without. We considered that each gene undergoing alternative splicing gave rise to three different isoforms with equal expression.

**Table 2.** Generating synthetic dataset 2

| Group | Total no. genes | Coverage | Genes without alternative splicing | Genes with alternative splicing | Transcripts with alternative splicing | Coverage for AS transcripts |
| --- | --- | --- | --- | --- | --- | --- |
| 1 | 5000 | 5 | 4500 | 500 | 1500 | 2 |
| 2 | 2000 | 50 | 1750 | 250 | 750 | 15 |
| 3 | 2000 | 100 | 1750 | 250 | 750 | 30 |

For simulation of PacBio reads, PBSIM parameters (read length, error probability by type, etc) were set to match those of dataset 5 containing reads of insert (see Supplement table 1). For simulation of MinION ONT reads, *PBSIM* parameters (read length, error probability by type etc.) were set to match those for MinION reads from a R9 chemistry dataset obtained from the Loman lab website (http://lab.loman.net/2016/07/30/nanopore-r9-data-release). Only 2d reads statistics were used.

### 2.3 Error correction

To test if the alignment results could be improved using error correction, the highest quality PacBio dataset (containing ROIs) was corrected. Error correction was performed using Racon (Vaser *et al*., 2017). Correction using Illumina reads, and self-correction were tested. Since self-corrected dataset proved to have better error profile, only this dataset was retained for the benchmark (Dataset statistics is given in Supplement table 1). Supplement table 1 displays error rate and read length statistics for all real datasets, including all datasets obtained using error correction.

### 2.4 Evaluated RNA-seq tools

We tested five RNA-seq alignment tools that have been updated recently reflecting that they are still being maintained.

#### STAR

Downloaded from https://github.com/alexdobin/STAR. Version 2.5.2b was used. STAR was run with parameters suggested at *Bioinfx study: Optimizing STAR aligner for Iso Seq data* from PacBio GitHub pages (https://github.com/PacificBiosciences/cDNA_primer/wiki/Bioinfx-study:-Optimizing-STAR-aligner-for-Iso-Seq-data), see Supplement Note 2.

#### Tophat2

Binaries were downloaded from https://ccb.jhu.edu/software/tophat/index.shtml and used with BowTie2. Version 2.1.1 was used, with default parameters for alignment. SAMTools version 1.2 were used to convert Tophat output from BAM to SAM format.

#### Hisat2

Binaries were downloaded from https://ccb.jhu.edu/software/hisat2/index.shtml. Version 2.0.4 was used, with default parameters for alignment.

#### BBMap

Downloaded from https://sourceforge.net/projects/bbmap/. The script mapPacBio.sh was used. BBMap version 35.92 was used. Reads were first converted to FASTA format (originally in FASTQ format) using samscripts tool (https://github.com/isovic/samscripts). The program was then run with the option fastareadlen set to a value appropriate for each dataset.

#### GMap

Source code was downloaded from http://research-pub.gene.com/gmap/. Version 2016-11-07 of GMap as used. GMap was used with default parameters, as recommended in the tutorial for using GMap with PacBio data (https://github.com/PacificBiosciences/cDNA_primer/wiki/Aligner-tutorial%3A-GMAP%2C-STAR%2C-BLAT%2C-and-BLASR).

### 2.5 RNAseqEval tool

Three of the five RNA-seq aligners were evaluated on resource usage and alignment quality. CPU and memory consumption were evaluated using a fork of the Cgmemtime tool (https://github.com/isovic/cgmemtime.git).

To evaluate the quality of each aligner, we developed RNAseqEval (https://github.com/kkrizanovic/RNAseqEval), meant to be a general tool for evaluating RNA-seq alignments. It is written in Python and contains two main scripts, one for evaluating data simulated using PBSIM and the other for evaluating real data or data whose origin is unknown. Both scripts require aligner output in SAM format which they compare to gene annotations and, in case of simulated data, alignment files in MAF format describing the origin of each simulated read.

#### 2.5.1 Evaluating synthetic data

The script for evaluating synthetic or simulated data currently works only on data simulated with PBSIM, but could be expanded in the future to support other simulators. Aside from aligner output in SAM format and gene annotations in GTF or BED format, the script takes a folder containing files generated by PBSIM. The folder containing PBSIM data needs to have a specific structure and follow a specific naming convention described in the program documentation.

For each read from aligner output, the script will use PBSIM generated MAF files and gene annotations to find its origin on the reference genome and will compare it to the alignment calculated by the aligner. The start and end position of an alignment and of read origin are compared, and an error of 5 nucleotides is tolerated. The script outputs summary information on how many reads were accurately aligned to their chromosome, strand and position of origin.

#### 2.5.2 Evaluating real data

The script for evaluating real data takes only aligner output in SAM format and gene annotation in GTF or BED format as its input. Because the origin of a read is unknown, the script will check annotations for genes with which the read overlaps, and then evaluate how well a read alignment matches exons and introns of that gene.

When matching beginning and end of an alignment to each exon in an annotation, an error of 5 nucleotides is tolerated. Similarly, an overlap between an alignment and an exon annotation needs to be at least 5 base-pairs to be considered valid.

## 3 Results

### 3.1 Baseline comparison

We first examined how alignment tools performed on the Illumina “baseline” dataset 0 (Table 3). We found that all aligners managed to align a large fraction of Illumina reads.

On datasets that include longer and more erroneous reads however (dataset 1 to dataset 8), there were large discrepancies across tools. In particular, Tophat2 and Hisat2 were unable to align any or hardly any read. Therefore, we did not consider these two tools in further analyses, and we focused on the remaining three aligners: BBMap, GMap and STAR.

**Table 1.**
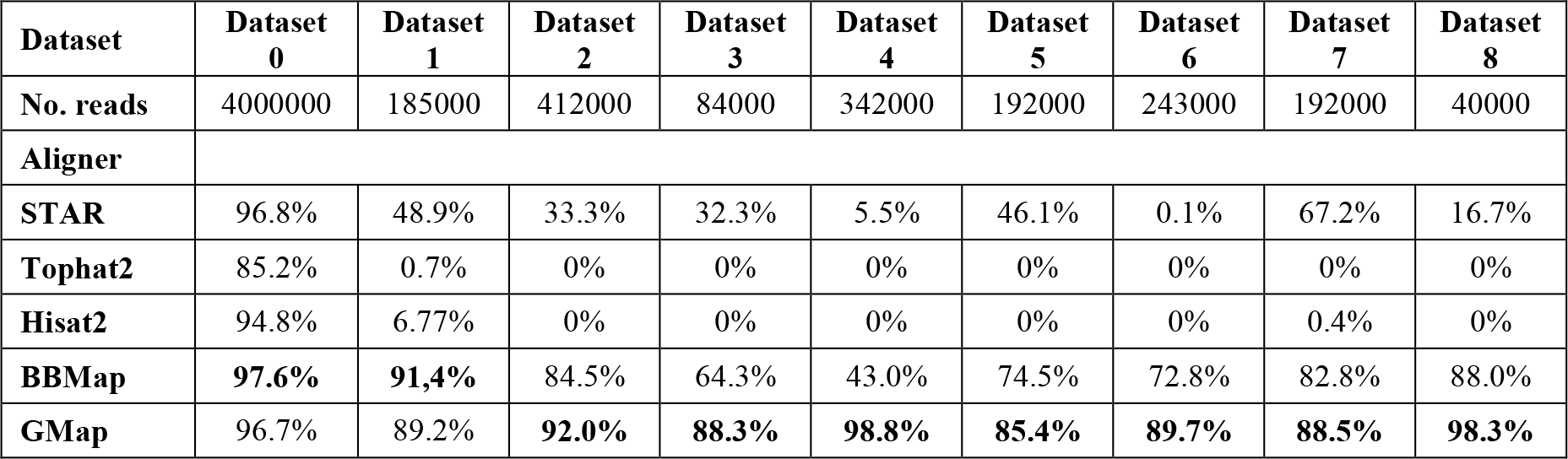
Percentage of reads aligned over all aligners and datasets

Based on the percentage of reads aligned, the best results were achieved by GMap, which aligned more than 85% of reads across the all tested datasets.

BBMap performed slightly better on Illumina (dataset0) and on synthetic *S. cerevisiae* PacBio dataset (dataset1, which contains very few multi-exon transcripts), but the fraction of reads aligned fell behind GMap on more complex synthetic datasets and real datasets (e.g., only 43% of the synthetic *H. sapiens* PacBio reads of dataset 4 were aligned).

STAR managed to align a large percentage of Illumina reads (96.8%), but its performance was uneven across third generation sequencing datasets, aligning from 0.1% to 67.2% of the reads, and often aligning less than half of the reads. STAR was seemingly affected by increased complexity of the datasets, as well as by increased error rates (Illumina and error-corrected PacBio datasets achieving the best performance).

Across all tools, error correction improved alignment rates, as can be seen from the comparison of dataset 5 and dataset 7.

In summary, for some aligners the percentage of alignment for third generation sequencing technologies reads was similar to what is achieved for Illumina reads. However, looking only at the number of the reads each tool managed to align to a genome is not a reliable measure of general alignment quality. For example, a tool could align most of the reads, but only on only a portion of their length, or it could align them at incorrect location on the genome.

### 3.2 Synthetic datasets

To get more insights into the quality of the alignments, we evaluated the aligners on 4 synthetic datasets generated from transcriptomes of varying complexity using the PBSIM tool (materials and methods), and supposed to reflect characteristics of the PacBio (datasets 1 to 3) and ONT MinION technologies (dataset 4). In these datasets, the precise origin of each read is known, allowing to assess the alignment quality by examining how well the alignment location matches the origin location in the genome. The alignment results for those datasets were evaluated using the RNAseqEval tool, as summarized in Figure 2.

**Figure 2.**
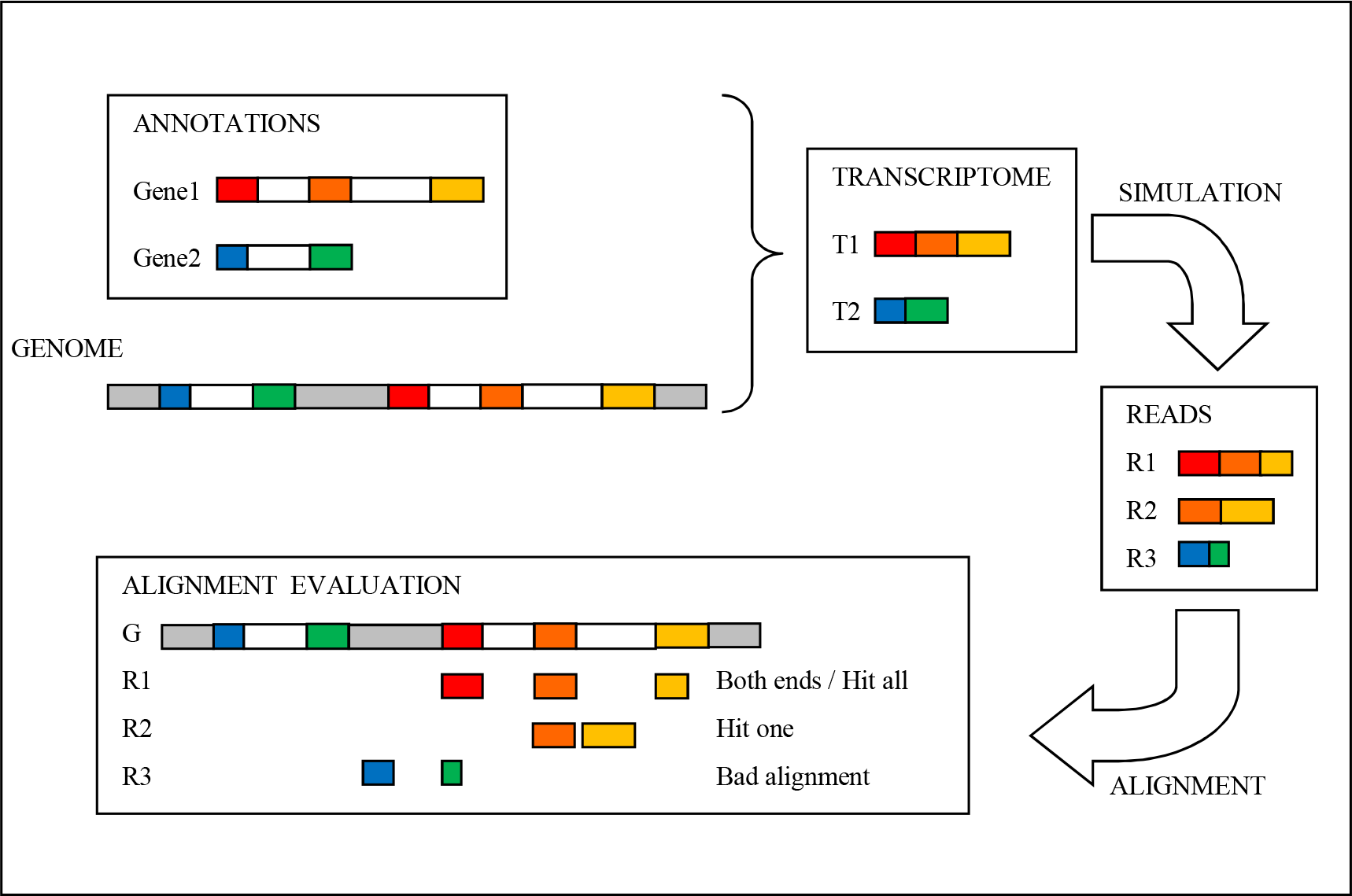
Simulating and evaluating synthetic datasets

Results of the evaluation on all synthetic reads are shown in Table 4. The evaluation on the subset of split reads (i.e., reads aligned to multiple non-contiguous locations on the reference genome) is also shown. Split reads, if aligned correctly, should overlap at least one exon-exon junction in the transcript of origin, and thus cover two or more exons. Percentage of reads shown in Table 4 are relative to the number of reads in input; the percentage relative to the number of aligned reads are shown in Supplement table 2.

Overall, the most accurate alignments were given by GMap and BBMap. In particular, GMap performed the best at aligning reads to correct general genomic locations, i.e., overlap exonic regions from which the simulated reads originated (hit all and hit one). BBMap however performed better at aligning the beginning and end of reads to their exact genomic position of origin (both ends), especially on lower complexity datasets.

Reads aligned by STAR mostly aligned to correct general genomic locations (hit all and hit one), and displayed very good match rates, however the low fraction of reads overall aligned (Table 3 and 4) did not allow this tool to compare favorably to GMap and BBmap. Moreover, STAR did not perform particularly well at correctly aligning the beginning and end of reads.

Datasets 2, 3 and 4 displayed a significant number of split reads, for example dataset 3 based on chromosome 19 of human genome included 60% of genes with alternative splicing. Focusing on split read statistics on those datasets, BBMap performed significantly worse than GMap and sometimes than STAR: on dataset 3 it managed to overlap all exons from a read origin (Split hit all) less precisely than STAR (10.2% Vs. 19.4%). For STAR, results for split reads were in line with its overall results, but the overall number of aligned reads being so low, STAR cannot be recommended for the alignment of third generation sequencing RNA-seq reads.

Overall, BBMap outperformed GMap in alignment precision on datasets with lower isoform diversity, but lagged behind in general alignment efficiency, sometimes by a large margin, on more complex datasets. This indicates that BBMap could be the best tool to cope with third generation sequencing error rate as a DNA-seq aligner, but should be used with caution to align split RNA-seq reads. In this setting, GMap shows the best performance and should be preferred, although the results on dataset 1 indicate that it is not the best tool to deal with high error rates of third generation sequencing data reads.

**Table 2.** Aligner evaluation on synthetic datasets. All results are displayed as the percentage of all reads in the dataset. The percentages of reads that were aligned is shown (without assessing the accuracy), the match rate of aligned reads, percentage of reads for which the beginning and the end are accurately placed (**both ends**), percentage of reads that overlap all exons of the read origin (**hit all**) and percentage of reads that overlap at least one exon of the read origin (**hit one**).

| Data set | Aligner | Aligned | Match rate | Both ends | Hit all | Hit one | Split reads | Both ends split | Split hit all | Split hit one |
| --- | --- | --- | --- | --- | --- | --- | --- | --- | --- | --- |
| 1 | STAR | 48.9% | **93.7%** | 21.6% | 46.5% | 47.1% | 1.89% | 0.8% | 1.19% | 1.78% |
|  | BMap | **91.4%** | 92.5% | **48.2%** | **87.0%** | **88.1%** | 3.46% | **1.0%** | **2.22%** | **3.30%** |
|  | GMap | 89.2% | 92.3% | 40.9% | 84.3% | 85.4% | 3.30% | 0.9% | 2.00% | 3.14% |
| 2 | STAR | 33.3% | **94.0%** | 4.1% | 27.7% | 30.7% | 23.9% | 1.4% | 19.3% | 22.3% |
|  | BMap | 84.5% | 89.9% | **10.5%** | 54.4% | 78.4% | 64.8% | **2.4%** | 36.7% | 60.7% |
|  | GMap | **92.0%** | 92.0% | 8.7% | **69.7%** | **85.4%** | 72.8% | 1.7% | **52.7%** | **68.5%** |
| 3 | STAR | 32.3% | **94.3%** | 3.9% | 27.5% | 30.5% | 23.1% | 1.7% | 19.4% | 22.4% |
|  | BMap | 64.3% | 86.2% | **10.0%** | 26.8% | 61.2% | 46.0% | 3.3% | 10.2% | 44.5% |
|  | GMap | **88.3%** | 91.8% | 8.0% | **64.8%** | **83.7%** | 70.0% | **4.4%** | **48.9%** | **68.0%** |
| 4 | STAR | 5.50% | 89.6% | 0.7% | 5.03% | 5.3% | 3.19% | 0.1% | 2.9% | 3.16% |
|  | BMap | 43.0% | 88.4% | 4.7% | 26.8% | 42.1% | 34.2% | 2.1% | 18.7% | 33.8% |
|  | GMap | **98.8%** | **90.5%** | **6.6%** | **83.6%** | **97.1%** | 80.4% | **3.0%** | **66.4%** | **79.8%** |

### 3.3 Real datasets

For real data, the origin of each read is not known, thus aligners were evaluated by comparing the read alignment locations to a given set of gene annotations. Some other relevant statistics, such as alignment match rate and number of expressed genes, were also extracted (Table 5). All real datasets consisted of technical replicates of RNA-seq on the same *D. melanogaster* sample sequenced on different platforms. Interestingly, these datasets were characterized by different error profiles (Supplement table 1).

**Table 5.** Aligner evaluation on real datasets. The table shows percentage of reads that were aligned (without assessing the accuracy), percentage of reads that overlap at least one exon (**exon hit**) and percentage of reads that overlap one or more exons in a sequence, corresponding to a gene annotation (**contiguous exon alignment**). All values are displayed as the percentage of all reads in the dataset. The table also shows the number of expressed genes and average match rate of aligned reads. Match rate is calculated as a percentage of aligned bases that are equal to the corresponding bases on the reference.

| Data set | Aligner | Aligned | Match rate | No. expressed genes | Exon hit | Contiguous exon alignment |
| --- | --- | --- | --- | --- | --- | --- |
| 5 | STAR | 46.1% | 92% | 8310 | 27.5% | 10.2% |
|  | BMap | 74.5% | 71% | 9371 | 45.1% | 20.1% |
|  | GMap | 85.4% | 88% | 10427 | 49.1% | 27.1% |
| 6 | STAR | 0.1% | 0.8% | 94 | 0.05% | 0.0% |
|  | BMap | 72.8% | 68% | 9166 | 45.3% | 15.5% |
|  | GMap | 90.1% | 82% | 10884 | 51.9% | 20.4% |
| 7 | STAR | 67.2% | 93% | 8515 | 39.0% | 18.6% |
|  | BMap | 82.8% | 72% | 9599 | 48.4% | 22.1% |
|  | GMap | 88.5% | 92% | 9985 | 50.4% | 31.4% |
| 8 | STAR | 16.8% | 83% | 1675 | 6.3% | 1.4% |
|  | BMap | 88.0% | 67% | 5449 | 36.3% | 12.2% |
|  | GMap | 98.3% | 81% | 5266 | 39.0% | 18.7% |

As expected from previous tests, GMap showed the best results, followed closely by BBMap. GMap was notably slightly better at aligning reads to annotated exonic locations in the genome. The match rate of aligned reads was roughly equal to the determined error profile for each dataset (Shown in Supplement table 1) thus suggesting that the reads are aligned to correct positions. GMap was even able to align ONT MinION data with a reasonable accuracy. It is interesting to note that by some criteria GMap shows better results on lesser quality dataset 6 (consisting of subreads) compared to higher quality dataset 5 (consisting of ROI) and dataset 7 (error corrected ROI).

Both BBMap and GMap reported a large percentage of ONT MinION reads aligned, however, match rate and exon hit percent-age were lower than for PacBio datasets, indicating that a larger percentage of those alignments were at an incorrect position.

STAR showed the worst alignment results. Reads successfully aligned displayed a high match rate, which might reflect the fact that STAR is unable to align reads with highest error rates, or that alignment settings are very conservative.

Supplement table 1 shows that error correction somewhat improved the error profile, increasing average match rate by 2-3 percent. However, even that slight improvement resulted in visibly better alignment results on dataset 7 for all aligners: more reads reported as aligned, more exons hit, more genes expressed and higher match rate. However, these results are achieved on significantly less reads. Considering that, the box plot results are consistent with those in table 4 and table 5.

Finally, we examined what fraction of the read length was aligned (Figure 3). The results are consistent with other measures of mapping quality, with STAR managing to align reads on a larger portion of their length compared to GMap.

**Figure 3.**
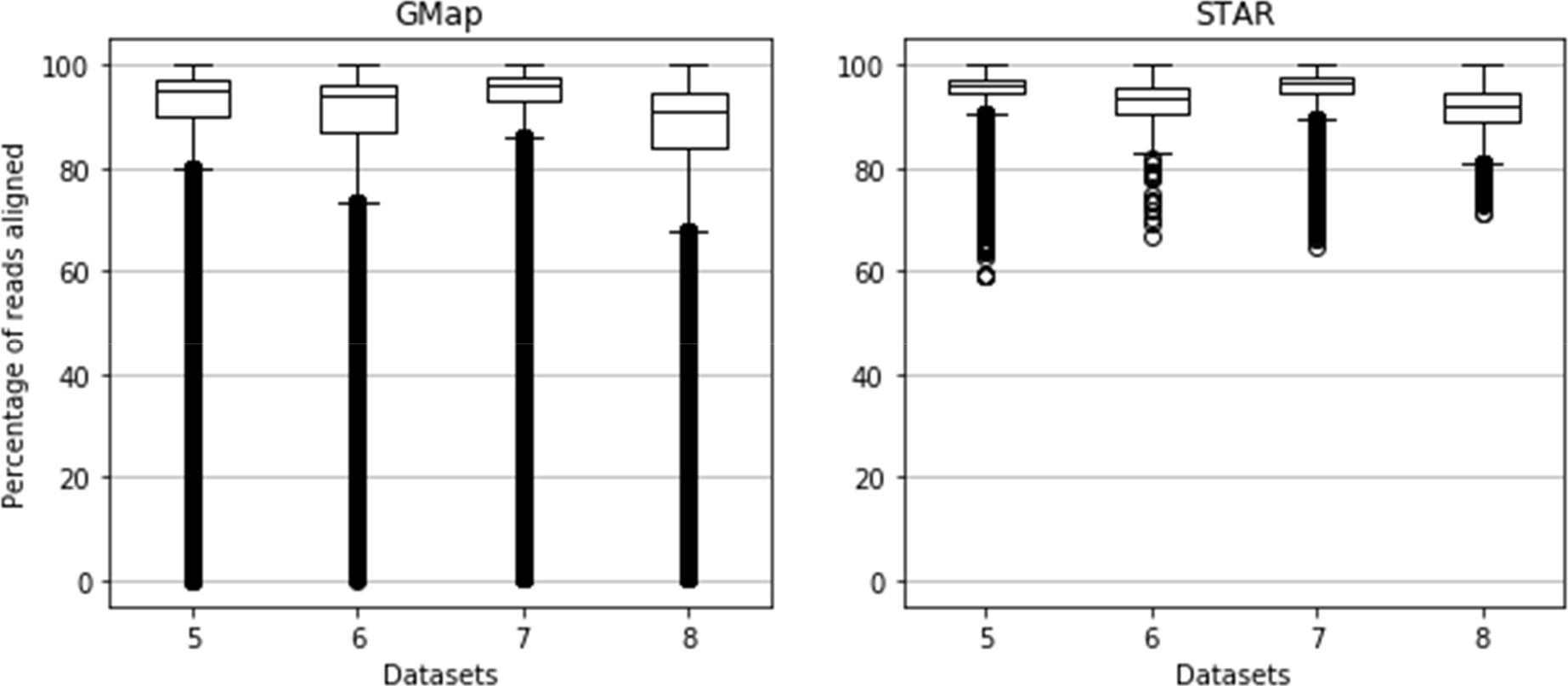
Box plot of aligned read percentage for STAR and GMap. Due to a large number of values, the plot contains many outliers.

BBMap results are not displayed because in the tested settings, all alignments are made on the whole length of the reads (global alignments). This behaviour has some implication in the reported results, as the alignment on both ends of the reads is sometimes incorrect, resulting in lower match rates. It could be a good idea to clip alignments resulting from BBMap, for example using the “local” flag, which converts global alignments into local alignments by clipping them if that results in higher scores. However, that option seems to cause exceptions in BBMap, resulting in terminated threads and in significantly lower number of aligned reads on real datasets. Because of that, we used results for BBMap global alignment, and do not show aligned read percentage box plot for BBMap.

### 3.4 Resource usage

To estimate the efficiency of each RNA aligner, CPU time and Maximum memory usage (Resident set size - RSS) were measured (Figure 4). Illumina data (dataset0) were omitted from this analysis because the focus of the paper is on third generation sequencing data.

**Figure 4.**
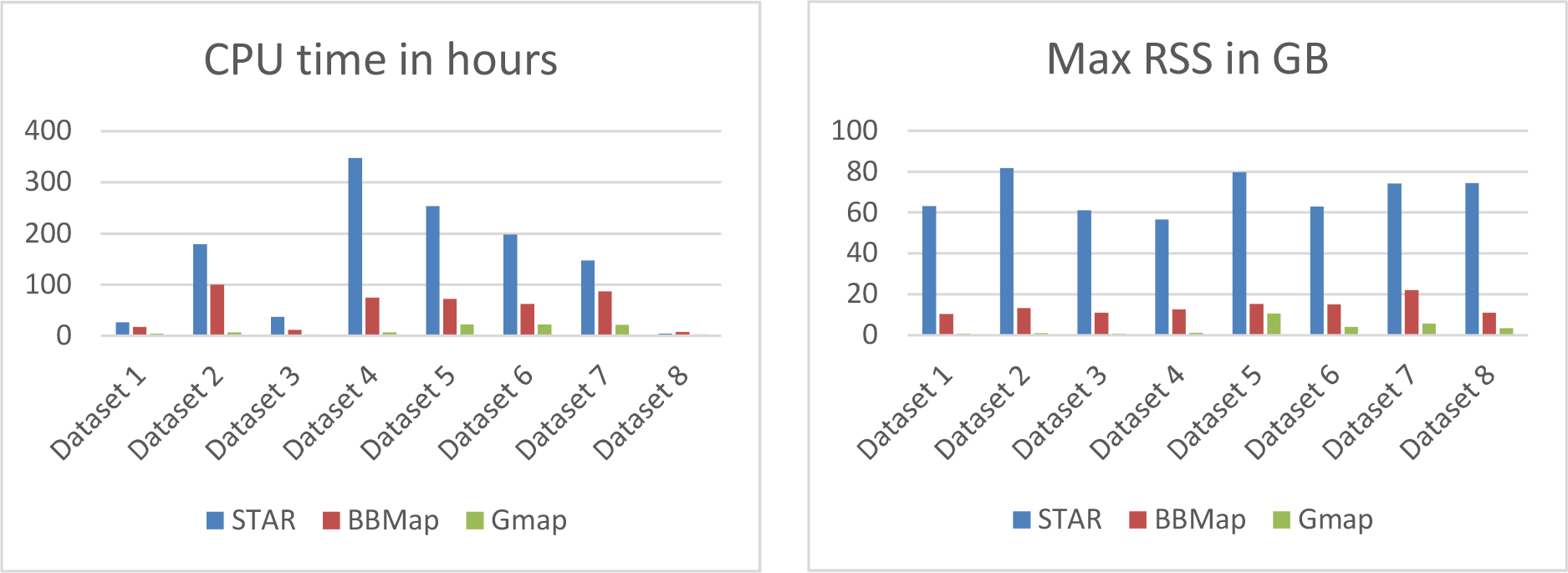
CPU time and memory usage

Running time seemed to depend on dataset size. In all settings, GMap used the least amount of memory and ran the fastest. STAR was the slowest and consistently used 60-80 GB of RAM. BBMap memory footprint was also consistently around 10-15 GB of RAM.

## 4 Conclusion

In recent years, third generation sequencing devices have been steadily establishing themselves in the area of genomic research. These technologies promise to solve problems caused by the short read length of the NGS. Regarding RNA-seq analysis, longer reads should notably improve transcript identification. However, third generation sequencing technologies also introduce new bioinformatics challenges, mostly due to their high error rate.

In this study we attempted to assess the ability of currently available RNA-seq alignment tools to work with third generation sequencing data. Five alignment tools were tested using real and synthetic datasets.

Hisat2 and Tophat2 were unable to align almost any read. STAR displayed only passable results on the least erroneous datasets, but failed almost completely on highly error-prone ONT MinION data.

BBMap, performed quite well, especially on PacBio ROI reads (which have lower error rates) and on simpler organisms with few multiexonic genes and low level of alternative splicing. This seems to indicate that although it is a splice-aware aligner, BBMap best performance is achieved on contiguous alignments (coming from DNA-seq for example), and might not be best suited for RNA-seq data.

Finally, GMap showed the best alignment results. It ran the fastest, used the least memory and usually produced the highest alignment rates, especially on complex datasets. This high rate of read alignment was sometimes at the cost of accuracy, and we observed that BBMap sometimes outperformed GMap in determining the correct beginning and end positions of aligned reads.

Overall, aligning third generation sequencing RNA reads is currently viable with some available tools, but we were surprised by the low precision on alignment location. Apart from dataset 1, no aligner attributed more than 10.5% of reads to their correct position of origin (+/-5 bases). It is not clear if this result is inherent to the high error rates of the technologies, or if it is due to alignment algorithms that were not originally developed for these types of data, or to the specific parameters used in this benchmark.

There is probably large room for improvement, by developing new more sophisticated and more sensitive algorithms, or by incorporating an error-correction step in bioinformatics pipeline before read alignment, since in our tests this visibly improved the alignment results.

